# The structure of a *C. neoformans* polysaccharide motif recognized by protective antibodies: A combined NMR and MD study

**DOI:** 10.1101/2023.09.06.556507

**Authors:** Audra A. Hargett, Hugo F. Azurmendi, Conor J. Crawford, Maggie P. Wear, Stefan Oscarson, Arturo Casadevall, Darόn I. Freedberg

## Abstract

*Cryptococcus neoformans* is a fungal pathogen responsible for cryptococcosis and cryptococcal meningitis. The *C. neoformans* capsular polysaccharide and shed exopolysaccharide functions both as a key virulence factor and to protect the fungal cell from phagocytosis. Currently, a glycoconjugate of these polysaccharides is being explored as a vaccine to protect against *C. neoformans* infection. In this combined NMR and MD study, experimentally determined NOEs and *J*-couplings support a structure of the synthetic decasaccharide, GXM10-Ac_3_, obtained by MD. GXM10-Ac_3_ was designed as an extension of glucuronoxylomannan (GXM) polysaccharide motif (M2) which is common in the clinically predominant serotype A strains and is recognized by protective forms of GXM-specific monoclonal antibodies. The M2 motif is characterized by a 6-residue α-mannan backbone repeating unit, consisting of a triad of α-(1→3)-mannoses, modified by β-(1→2)-xyloses on the first two mannoses and a β-(1→2)-glucuronic acid on the third mannose. The combined NMR and MD analyses reveal that GXM10-Ac_3_ adopts an extended structure, with xylose/glucuronic acid branches alternating sides along the α-mannan backbone. *O*-acetyl esters also alternate sides and are grouped in pairs. MD analysis of a twelve M2-repeating unit polymer supports the notion that the GXM10-Ac_3_ structure is uniformly represented throughout the polysaccharide. This experimentally consistent GXM model displays high flexibility while maintaining a structural identity, yielding new insights to further explore intermolecular interactions between polysaccharides, interactions with anti-GXM mAbs, and the cryptococcal polysaccharide architecture.

**Significance Statement:** This study utilized a combined NMR and MD approach to elucidate the structure of a *Cryptococcus neoformans* GXM synthetic decasaccharide (GXM10-Ac_3_), recognized by protective anti-GXM mAbs. The data revealed an extended structure in which the xylose/glucuronic acid branches and pairs of 6-*O*-acetyl esters predominantly alternate sides along the α-mannan backbone. MD analysis of a GXM polysaccharide predicts that the decasaccharide structure is uniformly represented in the polysaccharide. Additionally, the GXM exhibits high flexibility while maintaining structural identity. These findings lay the foundation for future studies aimed at understanding anti-GXM antibody-polysaccharide interactions.

## Introduction

*Cryptococcus neoformans* is a fungal pathogen with a worldwide distribution, that includes environmental and urban settings (1, 2), and is the causative agent of cryptococcosis and cryptococcal meningitis (3, 4). While most exposures do not lead to overt disease, *C. neoformans* can cause serious illness with a high mortality rate in immunocompromised individuals, especially in patients with HIV infection (5–7). Resistance to commonly used antifungal drugs is increasing, which creates a need for new therapeutics, e.g., vaccines (8).

*C. neoformans* is surrounded by a capsular polysaccharide (CPS) that is antiphagocytic and thus functions as a major virulence factor. Infection is accompanied by shedding of exopolysaccharide (EPS) which contributes to virulence by interfering with immune mechanisms. (9). One major component of both CPS and EPS is the glucuronoxylomannan (GXM) polysaccharide (PS). Currently an experimental *C. neoformans* GXM glycoconjugate vaccine is being investigated to combat this microbial infection (10). Glycoconjugates are extremely effective vaccines that reduce the incidence of infectious diseases caused by encapsulated bacteria. For example, licensed glycoconjugate vaccines utilizing the capsular PS of specific bacteria protect against *Haemophilus influenzae, Neisseria meningitidis, and Streptococcus pneumoniae* bacterial infections (11). We hope that a *C. neoformans* GXM PS will similarly result in a viable glycoconjugate vaccine protective against infection by this pathogen.

A glucuronoxylomannan (GXM) repeating unit (RU) consists of an α-(1→3) linked mannose (Man) triad backbone, modified by a β-(1→2) linked glucuronic acid (GlcA) branch on the 3^rd^ Man and β-(1→2)/β-(1→4) linked xylose (Xyl) branches (12). Additionally, Man O6 can be acetylated, but the degree of *O*-acetylation is strain-specific (13, 14). Unlike bacterial capsular PS, which have a regular RU, the *C. neoformans* GXM PS is heterogeneous and is classified into six motifs, differentiated by the location and number of Xyl branches (12, 15). These motifs were first characterized by nuclear magnetic resonance (NMR) of detergent extracted, de-*O*-acetylated, ^13^C-enriched native GXM polymer isolates. However, NMR signal degeneracy of the PS limited spectral interpretation to only the most abundant repeating motif units present in each isolate (16–22). Of the six motifs, the M2 motif, identified in Figure 1A, is commonly identified in *C. neoformans* serotype A, the most prevalent clinical form (23). The M2 motif triad is a six-residue RU consisting of a three-Man backbone with two branched β-(1→2) linked Xyl residues, followed by one β-(1→2) linked GlcA (14).

**Figure 1:**
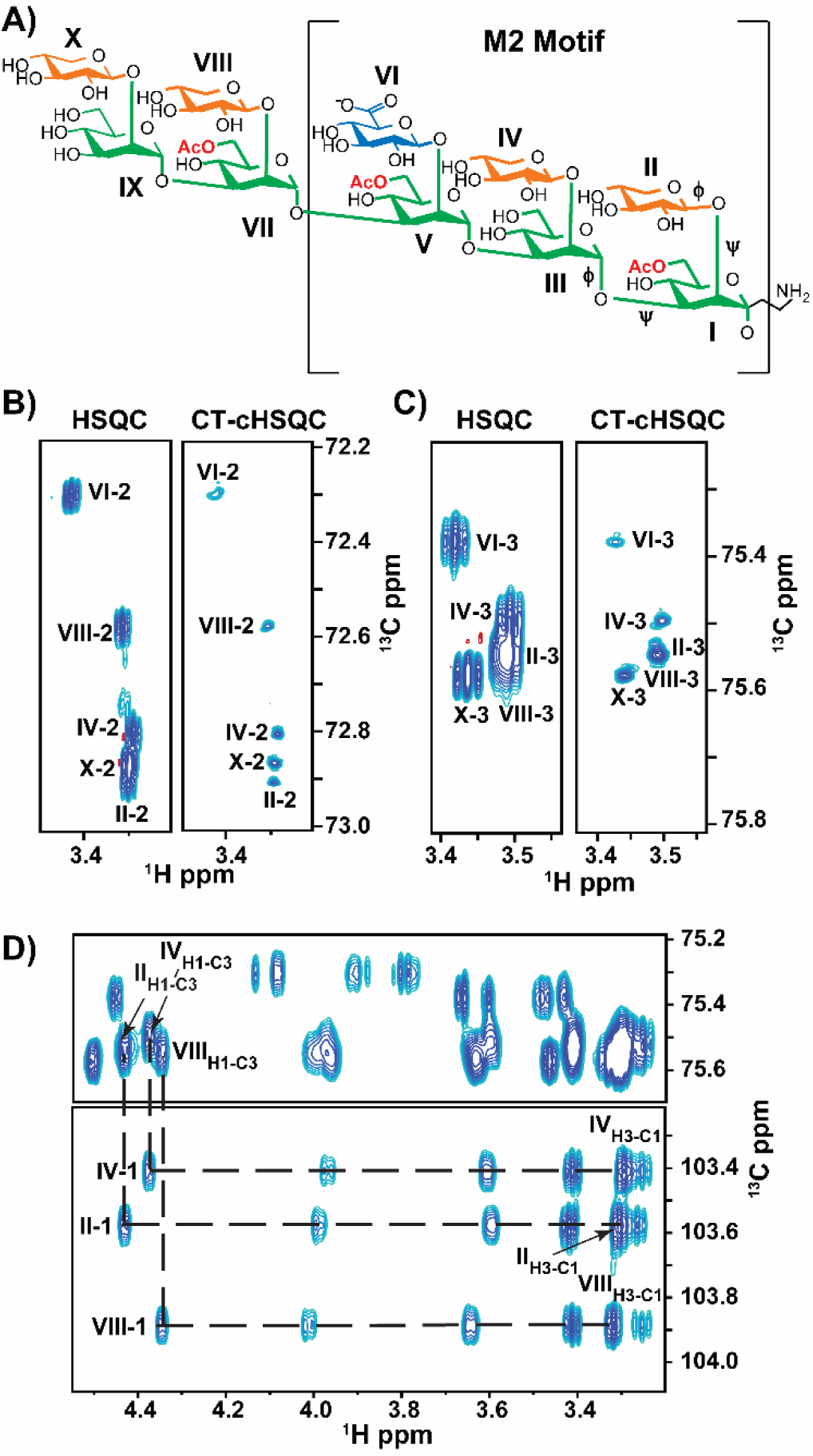
*C. neoformans* Glucuronoxylomannan (GXM) Synthetic Decasaccharide (GXM10-Ac_3_) resonances are resolved through ^1^H-^13^C CT-cHSQC and HSQC TOCSY experiments. (A) GXM10-Ac_3_ chemical structure contains the M2 motif, a common serotype A GXM repeating unit, and binds strongly to anti-GXM Abs (21). Glycans are color coded as follows: Man (green), Xyl (orange), and GlcA (blue). Transglycosidic φ and ψ torsions are denoted on the first three residues. **(B-C)** Comparison of resolutions achieved by a standard ^1^H-^13^C HSQC experiment relative to a CT-cHSQC experiment, with the latter allowing assignments of previously overlapping signals in a few hours for Xyl[II, IV, X]-2 **(B)** and Xyl[II,IV, VIII]-3 **(C)**. **(D)** ^1^H-^1^H crosspeaks between Xyl H1 and H3 obtained from a ^1^H-^13^C HSQC TOCSY experiment allows further differntiation between Xyl[II] and Xyl[III] H2C2 resonances.

Studies utilizing GXM as an antigen has yielded mixed results. The first cryptococcal glycoconjugate vaccine, a GXM-tetanus-toxoid (GXM-TT) glycoconjugate, elicited anti-GXM antibodies and was protective in mice (24, 25). While this first glycoconjugate vaccine was not pursued further, a similar GXM-TT vaccine was later shown to elicit antibodies with the same specificity as those produced from a cryptococcal infection in a murine model (26), some of which were protective against cryptococcal infection (27). Additionally, evidence showed that *O*-acetylation is critical for binding to protective antibodies (26, 28). While cryptococcal glycoconjugate vaccines can provide protection from infection, the PS utilized in the GXM-TT vaccine is a poorly characterized, heterogeneous mixture obtained from isolating *C. neoformans* PS. One strategy to develop a chemically defined glycoconjugate vaccine is the creation of a library of synthetic GXM oligosaccharides (OSs), which are then conjugated to carrier proteins. While synthetic tetra-to heptasaccharide conjugates did not bind to protective antibodies (29), a synthetic decasaccharide (GXM10-Ac_3_, Figure 1A) bound strongly to several protective antibodies (30).

To better understand Ab-PS interactions, we characterized the three-dimensional (3D) structural characterization of GXM10-Ac_3_ by high resolution solution NMR spectroscopy and molecular dynamics (MD) simulations and report these results herein. In this study, the NMR signal degeneracy is alleviated for GXM10-Ac_3_, which, together with modern NMR methods and higher fields to enhance signal resolution allowed a more thorough NMR characterization. To complement the NMR data, a GXM10-Ac_3_ 2 μs MD simulation was conducted and found to be in general agreement with the experimental NMR data. Once an experimentally consistent GXM10-Ac_3_ structure was derived, a GXM PS consisting of 12 M2 motif repeating units was studied by MD to test how well the combined experimental and theoretical decasaccharide results matched the extended GXM PS. The newly detailed structural characterization of the GXM PS provides insights into PS structural features which are likely to be important for recognition by the host immune system to neutralize the pathogen.

## Results

### NMR Measurements on GXM10-Ac_3_

The GXM10-Ac_3_ backbone is composed of five α-(1→3) linked Man, with β-(1→2) linked Xyl substitutions, except for the third Man, which bears a β-(1→2) linked GlcA. Additionally, Man I, V, and VII are acetylated at O6 (Figure 1A) (30).

To verify GXM10-Ac_3_’s primary structure and delineate connectivity, we collected ^1^H-^13^C HSQC (31), LR-HSQMBC (32) and HSQC-TOCSY (33, 34) at 700 MHz (^1^H frequency), 20 °C and pH 6.5. The standard ^1^H-^13^C HSQC (Figure S1) with 4.25 Hz/pt resolution was insufficient to resolve the Xyl residue peaks (II, IV, VIII, X) from each other without taking data for multiple days to obtain 0.5 Hz/pt resolution. Specifically, the H2-C2 resonances in residues II, IV and X (Figure 1B, HSQC) and H3-C3 in residues II, IV, and VIII (Figure 1C, HSQC) significantly overlap. We achieved full resolution of the H2-C2, H4-C4, and H5-C5 correlation peaks and improved resolution of the H3-C3 peaks by utilizing a CT-cHSQC (^1^H constant time, ^13^C-detected, ^1^H-^13^C HSQC) experiment, processed with SMILE-based linear prediction (35), in only four hours of data collection (Figure 1B, C, and Figure S2). However, as previously reported, this improved resolution comes with a reduction in sensitivity. Still, in the present study, the sensitivity in the shorter experiment was sufficient to make all assignments.

Residue-specific GXM10-Ac_3_ chemical shift assignments were obtained via a series of ^1^H-^13^C HSQC-TOCSY experiments using 30, 60, and 120 ms mixing times and confirmed with a LR-HSQMBC. Figure S1A shows a ^1^H-^13^C HSQC of the GXM10-Ac_3_ anomeric region. Each anomeric ^1^H-^13^C is unique and the ^1^H and ^13^C chemical shifts of each residue provide a “handle” from which the assignments could be made. For example, the ^1^H and ^13^C chemical shifts for the anomeric H1-C1 of residue II are 4.45 ppm and 102.48 ppm, respectively. In the 120 ms mixing time HSQC-TOCSY, five ^1^H cross peaks (3.99, 3.59, 3.42, 3.31, and 3.27 ppm) appear at a ^13^C chemical shift of 102.48 ppm. These ^1^H cross peaks correspond to all the ^1^Hs in the Xyl[II] ring (Figure 1D). Inter-residue linkage connectivity was determined via a LR-HSQBMC experiment. For example, the correlation peak at 4.19 ppm and 102.48 ppm connects the Man[I]-H2 with Xyl[II]-C1 (Figure S3). Table S1 provides the ^1^H and ^13^C chemical shift assignments for GXM10-Ac_3_.

In addition to the GXM10-Ac_3_ cross peaks, NMR spectra displayed peaks for partially de-*O*-acetylated decasaccharides. We assigned three additional decamers, each corresponding to the loss of one O-acetyl group at either Man[I]-C6, Man[V]-C6, or Man[VII]-C6 (Figure S4, Table S2). Assignments for the partially de-*O*-acetylated decamers differed measurably from GXM10-Ac_3_ only at the residue that lost *O*-acetylation and the proximal residues. These minor peaks were approximately 6-fold less intense than the fully *O*-acetylated decasaccharide. Complete chemical shift assignments of GXM10-Ac_3_ and its degradation products aided in the interpretation of the more complex NMR experiments used for structure determination (i.e., ^1^H-^1^H NOE and PIP-HSQMBC).

After assigning the chemical shifts, we used data from nuclear Overhauser effect spectroscopy (NOESY) (36) and ^1^H-^13^C pure in-phase HSQMBC (LR-PIPHSQMBC) (37) NMR experiments to help delineate the GXM10-Ac_3_ 3D conformations. ^13^C-edited NOESY spectra revealed 21 inter-residue ^1^H-^1^H NOEs (Table 1). Of the NOEs observed, one very weak peak with a signal to noise ratio (SNR) of 5 to 1 between Man[III]H3 and Xyl[II]H5 was detected. We did not observe any inter-residue NOEs between two branched glycans nor non-consecutive Man residues in our experiments, supporting a model of an extended structure with no persistent contacts between branched glycans. NOE signal intensities were classified as strong, medium, or weak, and this information was used to correlate with predicted NOEs from MD simulations (discussed below).

**Table 1.** Measured and predicted NOEs for GXM10-Ac_3_. NOE signals were classified based on their intensities as: S (strong); M (medium); or W (weak). H–H distance analysis from MD was done utilizing <R6> = < *r^−6^* >*^−1/6^*. Most H–H distances clustered around a single population, but in a few cases two populations were clearly distinguished. Cluster limits are indicated in the *D*_range_ column. All ranges are in bold, except minor populations (relative the main cluster) indicated with ‘normal’ font (two ranges in bold indicates similar populations).

| <sup>1</sup>H-<sup>1</sup>H NOE | NMR Obs. | MD $\langle R6 \rangle$ [Å] | MD $D_{\text{range}}$ [Å] |
| --- | --- | --- | --- |
| [I]H1 – [II]H1 | M | 2.5 | **1.8 – 2.4 / 2.4 – 4.0** |
| [I]H1 – [II]H5A | W | 4.8 | **3.6 – ~6.0** |
| [I]H1 – [II]H5E | W | 5.9 | **4.8 – <6.0** |
| [I]H2 – [II]H1 | S | 2.4 | **1.9 – 3.2** |
| [I]H2 – [III]H5 | W | 2.5 | **1.9 – 3.5** |
| [I]H3 – [III]H1 | S | 2.5 | **2.0 – 3.0** |
| [III]H1 – [IV]H1 | M | 2.4 | **1.8 – 2.4 / 2.4 – 4.4** |
| [III]H2 – [IV]H1 | S | 2.4 | **1.9 – 3.2** |
| [III]H2 – [IV]H2 | W | 3.8 | **4.1 – 4.7** |
| [III]H3 – [II]H5A | W | 3.5 | **2.7 – 4.7** |
| [III]H3 – [II]H5E | W | 2.7 | **2.0 – 3.6 / 3.6 – <6.0** |
| [III]H3 – [V]H1 | S | 2.4 | **1.9 – 3.0** |
| [V]H1 – [IV]H1 | M | 2.2 | **1.7 – 3.1 / 3.7 – 4.5** |
| [V]H2 – [IV]H1 | S | 2.5 | **2.0 – 3.5** |
| [V]H2 – [VII]H1 | S | 2.3 | **1.8 – 2.9** |
| [VII]H1 – [VIII]H1 | S | 2.3 | **1.7 – 3.6** |
| [VII]H2 – [VIII]H1 | S | 2.5 | **1.9 – 3.5** |
| [VII]H2 – [IX]H3 | W | 4.4 | **3.6 – 5.2** |
| [VII]H3 – [IX]H1 | S | 2.3 | **1.9 – 2.9** |
| [IX]H1 – [X]H1 | M | 2.4 | **1.7 – 2.8 / 2.8 – 4.2** |
| [IX]H2 – [X]H1 | S | 2.5 | **1.9 – 3.3** |

^1^H-^13^C PIP-HSQMBC NMR experiments provided transglycosidic ^3^*J*_CH_ values, which we substituted into parameterized Karplus relations to obtain phi (φ) and psi (ψ) transglycosidic torsion angles (Table S4). The φ and ψ torsions derived from transglycosidic ^3^*J*_CH_ values were similar for most cases, ranging from 2.5 Hz to 4.2 Hz for φ and 3.4 Hz to 5.5 Hz for ψ torsions. The ^3^*J*_CH_ for H-C-O-C Man[I]-Man[III] and H-C-O-C Man[III]-Man[V] were undetectable using the conditions in our experiments. While each transglycosidic ^3^*J*_CH_ value yields four potential torsion values, we narrowed down these possibilities to the most likely torsions by incorporating the results from MD analysis, as discussed next.

### Molecular Dynamics Simulation of GXM10-Ac_3_

Previous MD results on a family of GXM and related oligomers reported, that “the mannan backbone was consistently extended and relatively inflexible” (38). Our study used a different force field and a significantly extended duration of MD trajectories (from 100 ns to 2200 ns), nonetheless yielding a similar result regarding the mannan backbone average torsion values (Figure S5). However, the dynamic behavior of the OS displayed in our present, extended study, seems to be better described as flexible, given the range of torsion values visited through the MD trajectory. The torsion maps derived for only the penta-mannose backbone and GXM10 (w/wo Ac_3_) demonstrate that the duration of the MD was long enough for all potential conformations to be fully explored, and that the effect of adding substituents to the backbone is, as previously reported by Kuttel et al., mainly limited to narrowing the spread of torsion angle values reachable (Figure S5).

To obtain the most reliable structural model of GXM10-Ac_3_, we compared experimental NMR data with MD derived results. Observed NOEs are usually indicative of hydrogen atoms located at average distances less than 5 Å, however, motion often results in reduced NOE peak intensities, making them appear to be farther apart than they are in a model. We predicted NOE’s from the MD trajectory with a cutoff of 5.0 Å (see materials and methods for details). Out of 21 experimentally detected NOEs (Table 1), 19 H-H distances calculated from the MD trajectory (90%) agree with the ^1^H-^1^H NOEs obtained. Table 1 contains the range of distances derived from the trajectory, which reflects the weighted NOE distance averages (<*r*^6^>)^−1/6^. In most cases, a single range suffices to describe the MD derived distances, but in five cases two populations were clearly distinguished for which ranges are reported. These cases can explain some of the minor differences between observed and predicted NOEs. In addition, two NOE signals were weak as compared to the weighted distances, 2.5 and 2.7 Å, calculated from MD. These NOEs correspond to distances associated with Man[I] and its linked Xyl[II] to the contiguous backbone Man[III]. These discrepancies are most likely explained by a more flexible Man[I] (and its branched Xyl[II]) than predicted by MD. Overall, the excellent agreement with experimental NMR data lends support to the structures obtained from the MD. Furthermore, consistent with experiment, no NOEs between branching residues were predicted from the MD results.

Next, we analyzed the MD trajectory and plotted the low energy regions on a (φ, ψ) transglycosidic torsion angle map (Figure 2). The GXM10-Ac_3_ Man backbone glycosidic torsions show low energy conformations (< 2 kcal/mol) for a relatively narrow distribution of φ torsions (+/− 15°) and a wider distribution of ψ torsions (~ +/− 30°, Figure 2A). These values indicate a limited, but still broad, range of motion for the Man backbone of the GXM10-Ac_3_. Still broader is the Man-Xyl transglycosidic torsion angle population, with the presence of two or more local energy minima (Figure 2B.). Nevertheless, a single preferred low energy region is distinguishable in all cases with populations one or two orders of magnitude larger than for alternative regions. Similarly, the Man[V]-GlcA[VI] transglycosidic torsion shows a preferred low energy region with the addition of a distinct small secondary region centered at ~(15°, −65°) (Figure 2C).

**Figure 2.**
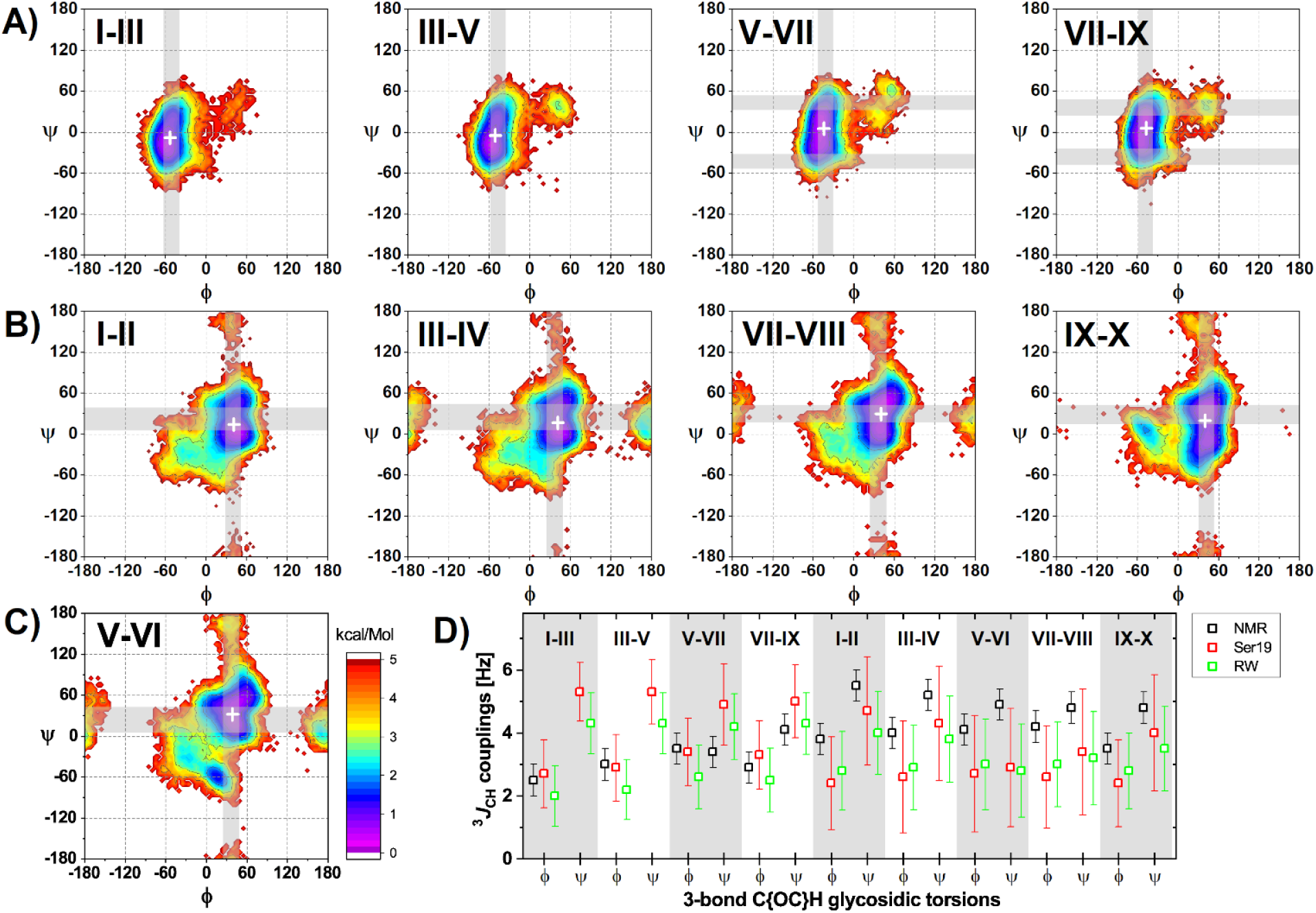
GXM10-Ac_3_ Transglycosidic torsion and ^3^*J*_CH_-couplings analysis. **(A-C)** Torsion energy heatmaps generated from the MD trajectory. The most likely predicted (low-energy) conformations are denoted in purple/blue (< 2 kcal/mol). Gray bands correspond to predicted torsion values from experimental ^3^*J*_CH_ NMR values utilizing the Karplus equation as parameterized in Reeves and Wang 2022 (40) which best agree with average MD trajectory torsions (white +). Table S3 further details all calculated torsions. **(A)** Results for Man-Man backbone linkages. **(B)** Energy maps for Xyl-Man branched linkages. **(C)** Analysis for the GlcA-Man branch linkage. **(D)** Comparison of measured ^3^*J*_CH_ values to MD predicted mean values using two recent approaches (see Methods).

Next, we compared the experimental transglycosidic ^3^*J*_CH_’s values to the averaged torsions from MD simulations (Figure 2D, Table S3 and S4). While *J*-couplings predicted from MD simulations are heavily influenced by higher populated regions, gross departures from experimental values would indicate problems with the simulation. Figure 2D shows experimental and predicted ^3^*J*_CH_ utilizing two recently published approaches (39, 40). Considering experimental errors of +/− 0.5 Hz and resulting MD standard deviations (+/− 1-2 Hz) both sets of values agree. This is important in that the MD results allow disambiguation of torsion values from experimental couplings, which are the result of conformational averaging. Moreover, predicted φ torsions mirror the experimental ones. For the measured ψ-associated couplings, the result is more ambiguous due to both the range of couplings obtained and the broad distribution of torsions predicted by MD. In Figures 2A-C, we show calculated torsion values from experimental *J*-couplings as gray bands overlayed to the energy maps. For clarity, we did not include torsion regions unlikely to contribute to the measured *J*-couplings based on the MD analysis. Importantly, Figure 2 shows that no inconsistency is observed between the four experimentally compatible, ^3^*J*-coupling derived torsion angles and the low energy regions.

### GXM10-Ac_3_ Structural Model

The next goal was to identify, if possible, a representative model from the trajectory for GXM10-Ac_3_ in solution. As a first approximation to this goal, we calculated a 2D-RMSD map of 2200 MD frames from a 2.2 μs trajectory (hence, 1 frame/ns), selecting the 60 ring atoms of the decasaccharide (10×6 = 60 atoms). The map shows (Figure S6) that GXM10-Ac_3_ stays in a similar shape compared to the initial conformation for about 75 ns, after which it switches to a new conformation in which it remains through the conclusion of the simulation, except for a couple of short-lived departures. Thus, we restricted the production analysis to 2 μs starting at the 100 ns mark. Taking 2000 uniformly sampled structures (one per ns), we performed RMSD minimization on the 60 ring atoms of the first frame of the 2000 set and obtained the resulting average structure (Figure 3A and supporting movie S1). The resulting averaged GXM10-Ac_3_ conforms closely enough to an energy minimized structure for the backbone Mans and three of the branched Xyls [II, IV, and VII]. GlcA[VI] and Xyl[X] are the rings more distorted relative to regular pyranose rings, but still clearly recognizable (6-ring atoms RMSD to minimized GlcA and Xyl pyranose rings were 0.28 Å and 0.31 Å, respectively). This is truly a remarkable result, as the branched glycans have ample freedom to visit a broad range of different orientations, as illustrated in Figure 3B, where 500 models (sampling 500 ns of the trajectory) of the Man6Ac[V]-GlcA[VI] branch were overlayed with RMSD minimization on the six ring atoms of Man[V]. The orientation of the branch was chosen to have an unobstructed view of the β-(1→2) glycosidic linkage and the corresponding (φ, ψ) torsions. Two populations can be distinguished, with an approximate ratio of 95% and 5% respectively. Naturally, the major population referred here correspond to the large purple/blue region in Figure 2C. The acetyl group [V] is also depicted to show its almost total freedom of orientation. Supporting movie S2 shows the dynamic behavior of GlcA for these 500 frames, utilizing the *smooth* Pymol script (with default values) for clarity. Similar results are obtained for all branches.

**Figure 3.**
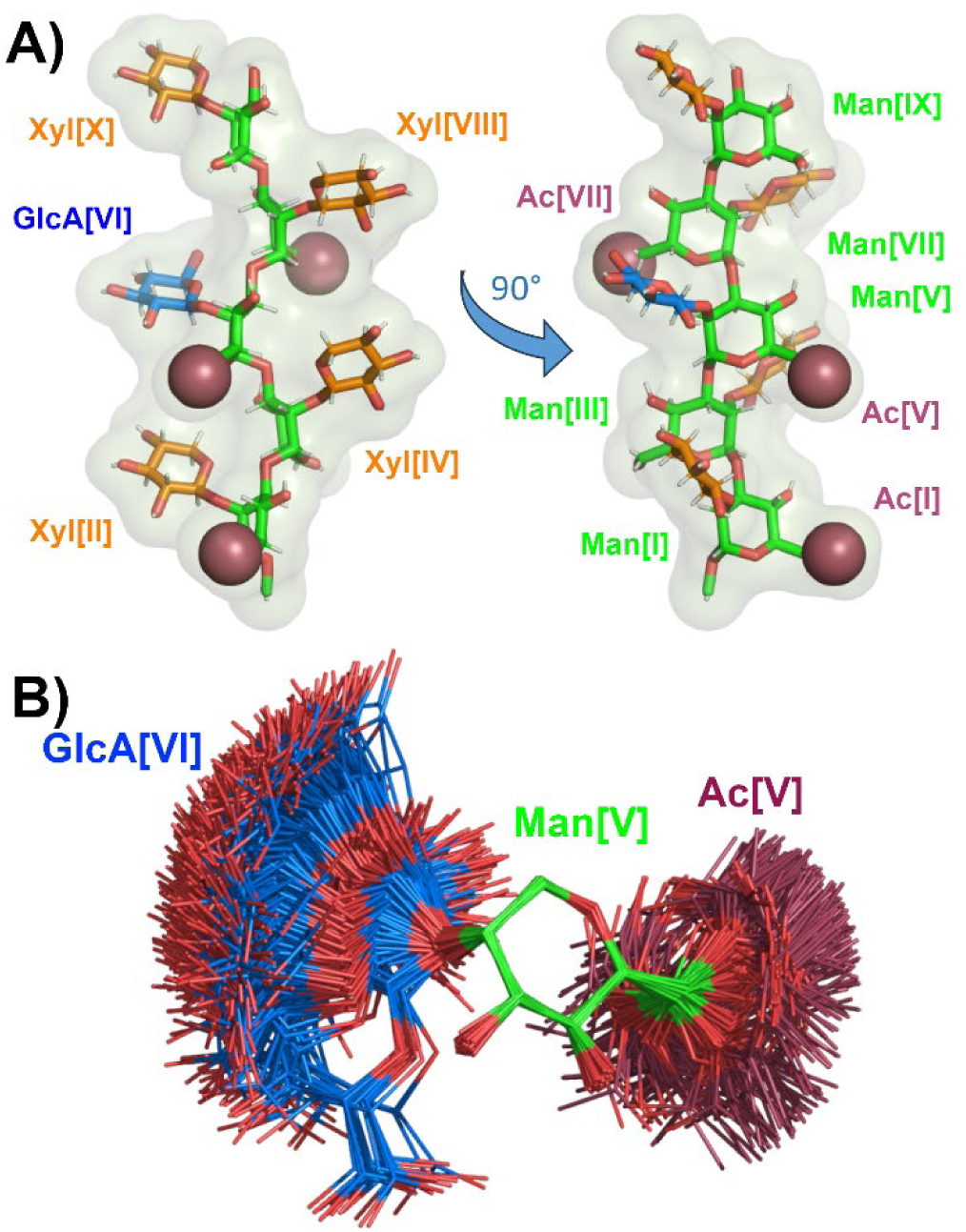
GXM10-Ac_3_ Structural Model. **(A)** The averaged MD structure of GXM10-Ac_3_ (GXM10ref) is consistent with NMR derived experimental parameters. The resulting model has an extended Man backbone with each contiguous Man in an antiparallel orientation (green), which results in the Xyl (orange) and GlcA (blue) branches alternating sides alongside the Man backbone. The Man[I] and Man[V] *O*-acetyl groups (raspberry) align on the same side of the Man backbone while Man[VII] 6Ac and GlcA carboxyl side chain align approximately on the opposite side. **(B)** Spread of orientations of the GlcA branch relative to the Man to which is linked (500 overlayed structures from the last 500 ns MD trajectory of GXM10-Ac_3_), showing broad freedom of movement which fluctuates between a “face-on” and “edge-on” orientation. Similar results are obtained for all branches.

Using the averaged structure in Figure 3A as reference (referred to as GXM10ref onward), we conducted a more granular analysis calculating the RMSD against GXM10ref for models sampled every 40 ps (50,000 frames in 2 μs), which resulted in overall RMSD (60 ring atoms) of 1.6 ± 0.6 Å. The frame along the trajectory with the lowest RMSD (0.56 Å) is overlaid to GXM10ref in Figure S7 to provide a concrete example of how RMSD values and conformations are related. The main contributions to the RMSD value in this example can be summarized by Xyl[VIII] being misoriented by ~90° and Xyl[II, X] slightly displaced.

Thus, calculations predict, and NMR results support, the Man backbone as extended with Man-Man transglycosidic torsions fluctuating about central φ and ψ values. Interestingly, the average consecutive Man residues alternate orientations as result of the α-(1→3) linkage, since the αC1 – O3’ bond (prime indicating the next Man residue) is axial and the O3’ – C3’ is equatorial, and so forth (this is most easily appreciated from the left model in Figure 3A). Thus, all axial bonds in contiguous Man residues are antiparallel including the substitution bonds Man[C2-O2]. Consequently, the Xyl/GlcA branches also alternate sides of the Man backbone, as shown in Figure 3A. Moreover, the Man[I] and Man[V] *O*-acetyl groups are on the same side of the mannan backbone and close in space, while 6Ac[VII] is located on the opposite side. As shown in Figure 3B (and accompanying movie S2), the GlcA branch has one main population which fluctuates between two extreme orientations associated to ψ ≳ 60° (edge-on, where the plane of the ring is approximately perpendicular to the Man backbone) and ψ ≲ 20° (face-on, where the plane of the ring is parallel to the Man backbone), hence, GXM10ref in Figure 3A presents an in-between orientation. No significant energy barrier seems to exist between these different shapes. The energy distribution displayed in Figure 2 arise from steric interactions of the salient carboxyl group in GlcA which preferably orients the residue with the ring mostly parallel to the neighboring Xyl[II, X]. The alternative 5% minor conformation has the plane of the ring oriented perpendicular and off-line the xylose residues.

### GXM Polysaccharide Molecular Dynamics Simulation

To structurally characterize the extended PS, we performed a 2 μs MD simulation on a 12 M2 motif RU GXM acetylated PS; thus, with the backbone consisting of a 36 α-(1→3) linked Man chain. The previous study reported was limited to 6 RU and 500 ns (38). Visual inspection, distance analysis, H bonds, and torsion angles didn’t reveal any pattern that could indicate the persistence of multi-residue structural features beyond what was previously described. Nevertheless, we decided to compare the GXM trajectory to GXM10ref, described in the previous section, over the length of the PS, which for a 12 RU contains 11 such segments centered at GlcA residues, as illustrated in Figure 4A. The resulting RMSD values, for each GXM segment (on 60 ring atoms) over the 2 μs trajectory, are shown in panel 4B. For clarity, RMSD instantaneous values were averaged over a running window of 20 ns, otherwise the wide variation of consecutive points obscures the ability to discern any trends over the trajectory. As can be appreciated, most GXM10-Ac_3_ segments start in arbitrary initial conformations not resembling that of the reference model (RMSDs > 2 Å), but by approximately 1 μs all segments have found their way to a reference model-like shape with an RMSD value of ~1.3 Å, in which they mostly remain for the duration of the simulation. This result supports the conclusion that the experimentally consistent structure GXM10ref displayed in Figure 3A is uniformly present along the GXM PS.

**Figure 4:**
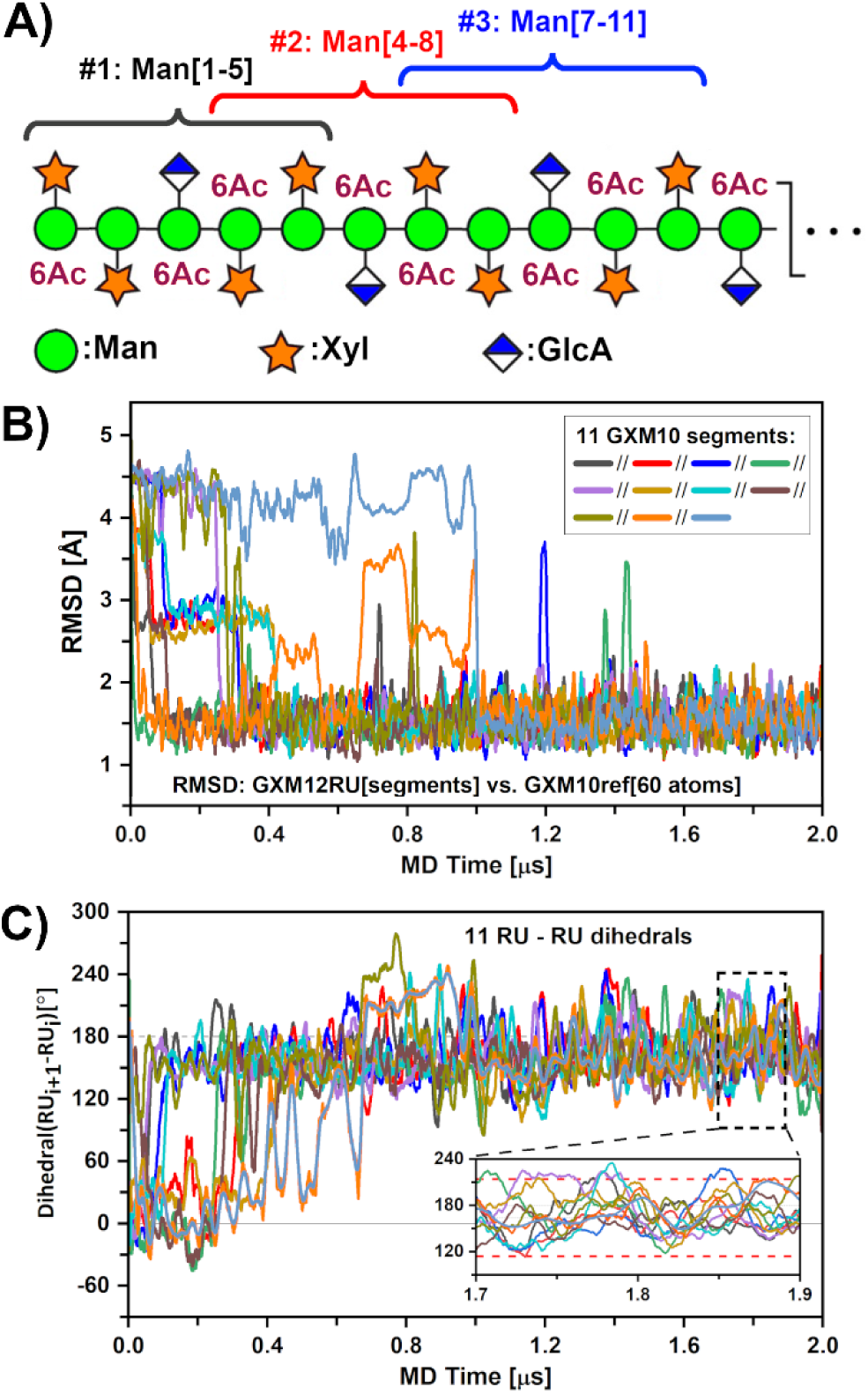
GXM-12RU (M2 motif) polysaccharide MD characterization. **(A)** Scheme of GXM12-RU PS first four repeating units (CFG notation), highlighting overlapping GXM10-Ac_3_ units, assessed over a 2 μs MD trajectory. **(B)** RMSD of PS segments against GXM10ref. After ~1.5 μs, all segments converged to a low RMSD ~1.5 Å. **(C)** Analysis of the rotation pitch of the RUs, using dihedrals between consecutive GlcA glycans (represented by Man{C2}-GlcA{C4} vectors). Averaged values converge to ~164°±50° toward the end of the simulation (indicated on inset with red dashed lines). In both (B) and (C) curves were smoothed using 20 ns running windows average for clarity.

Next, and given the segmental persistence along the PS of a preferred shape, we investigated whether the extended PS structure could propagate into a helical pattern. To do this, we chose to define a vector originating in the backbone and ending on GlcA. Thus, we defined vectors from Man{GlcA}-C2 to GlcA-C4 for each segment and measured all dihedral angles between consecutive RUs (Figure S8), following their evolution throughout the trajectory (Figure 4C). Utilizing a 20 ns running average reveals that the PS settles in a steady pattern after ~1.1 μs with dihedral values around 160°. A precise calculation from the representative portion shown in Figure 4C inset using instant values, results in an overall dihedral distribution of 164° +/− 50°. On average the PS completes one turn every 2.2 RU (~360°/164° = 2.2). Moreover, this tendency is easily reversed based on the large SD of ± 50° from instant dihedral values. Since the pitch of the helix is much larger than GXM10Ac_3_ (1.6 RU) which is the smallest oligosaccharide unit that binds strongly to many anti-GXM mAbs (30), it is unlikely that the helix is important for Ab recognition for these mAbs. However, in a recent study of synthetic GXMs a longer oligosaccharide consisting of 2.6 RUs was shown to bind stronger than GXM10Ac_3_ for at least two mAb 3E5 IgG_3_ and IgA (53). Thus, while this structural element may not be important for all anti-GXM mAbs, the ability of the PS to form a transient helix could be important for some anti-GXM mAb recognition and further experimental evidence is necessary to address this. Supporting movie S3 illustrates the dynamic behavior of the PS over the last 500 ns of the MD trajectory.

We finalize this work with Figure 5, showing three views of 6 RUs (out of 12) from the last MD trajectory frame to illustrate the GXM PS structural characteristics. This can be better described as an extended Man backbone, with alternating orientations, decorated with noticeable pairs of *O*-acetyl moieties which switch sides over the length of the PS. Additionally, the GlcA carboxylic acid group in each RU appears near the 6Ac pair of the following RU. We have used sphere representations to easily locate both 6Ac groups and the oxygen atoms in the carboxylic acid moieties in GlcA. The bulky Xyl/GlcA, alternate sides and add volume to the chain but with considerable orientational freedom. Therefore, the PS appears to maintain steady structural features while retaining ample liberty to depart from the resulting average values. The GlcA and *O*-acetyl groups placement at defined intervals alternating orientation suggest the hypothesis, which we will investigate in another report, that PS molecules interact through Ca^2+^ bridges (41) to create a complex network from which the capsule emerges.

**Figure 5.**
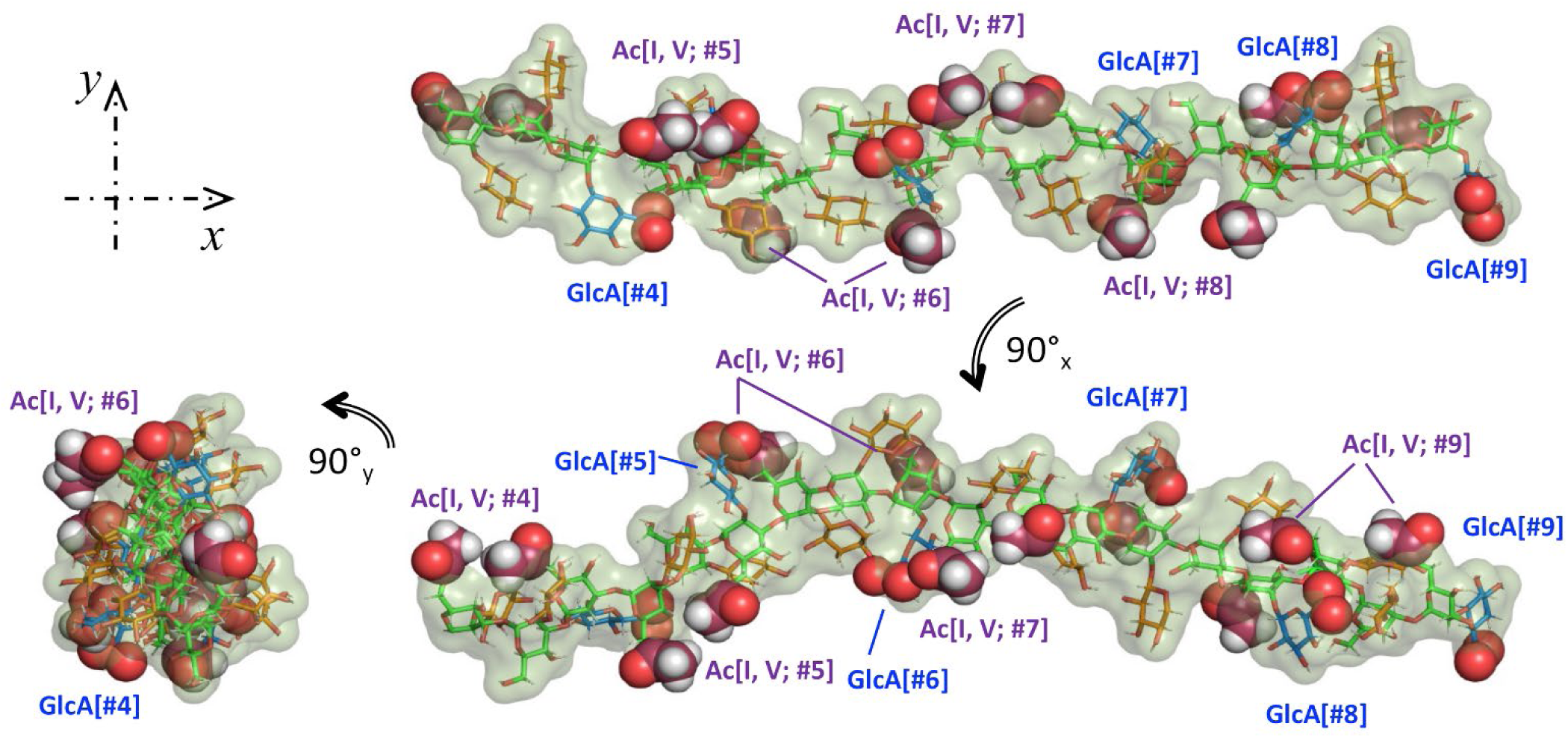
Three views of 6RU (#4-#9) of the last frame of the GXM PS MD. Color scheme: green(Man{C}); red(Xyl{C}; blue(GlcA{C}; raspberry(GlcA{O6A,B}); orange(6Ac{C}). Spheres representation used for 6Ac and Glc{O6A,B) } to facility identification.

## Conclusions

Until now a *C. neoformans* GXM PS structural characterization has remained elusive because isolated materials have been large and heterogenous. Hence, we focused instead on characterizing the structure of an immunologically active synthetic GXM10-Ac_3_ oligosaccharide utilizing modern NMR techniques combined with extended MD simulation data. Excellent agreement between NMR and MD structural analysis provides strong support for the derived representative model presented in Figure 3A, allowing us to draw general conclusions as to the important features regarding Ab recognition. Because the regularity of the Man backbone with rings alternating orientation, consecutive Xyl/GlcA branches are found on alternating sides of the PS. As a result, every other branched residue is close in space but with enough separation to allow ample orientational freedom. This also applies to the *O*-acetyl groups, with Man[I] and Man[V] *O*-acetyls close in space and Man[VII] *O*-acetyl on opposite sides of the Man backbone. The GXM 12RU PS MD experiments allowed us to generalize these observations to the PS, showing that the experimentally verified GXM10-Ac_3_ shape is, on average, uniformly represented in the PS. In addition, MD predicts that although the PS tends to form a helical pattern the pitch is significantly longer than GXM10-Ac_3_. Thus, our data suggest that the structural elements important for Ab recognition are the proximal pair of *O-*acetyl groups. Overall, the deduced GXM PS structure provides a foundation for future studies to characterize interactions between cryptococcal GXM PS and host immune factors.

## Materials and Methods

### Synthesis of the Decasaccharide

Synthesis of the decasaccharide was reported previously (30), which used a convergent building block approach, utilizing di-and tetrasaccharide thioglycoside building blocks (42, 43). The global deprotection were then carried out using optimized conditions (44, 45).

### NMR Analysis of Synthetic Decasaccharide

All NMR experiments were performed at 700 MHz ^1^H frequency using a Bruker Avance III HD console and 5 mm triple resonance xyz gradient cryoprobe at 20 °C. 2.5 mg of the GX10-Ac_3_ synthetic decasaccharide were dissolved in 300 µL of in 20 mM phosphate buffer at pH* 6.4 with added DSS as an internal reference in ~99% D_2_O. Two-dimensional NMR spectra were acquired using Bruker 3.6.5 software (http://www.bruker.com). Data were collected with a ^1^H spectral window of 7002.80 Hz (10 ppm) and a ^13^C window of 8802.92 Hz (50 ppm) with carrier frequencies of 4.705 ppm and 83.00 ppm in ^1^H and ^13^C, respectively. For ^1^H-^13^C HSQC (Bruker pulse sequence hsqcedetgp), HSQC-TOCSY (Bruker pulse sequence hsqcdietgpsi) experiments, 2048 points were taken in ^1^H and ^13^C. In the LR-HSQMBC (Bruker pulse sequence hsqcetgpipjcsp2), PIP-HSQMBC (37), and NOESY (Bruker pulse sequence hsqcetf3gpno) experiments 4096 and 2048 points were taken in ^1^H and ^13^C, respectively. ^1^H NOE strength was assigned based off the NOE crosspeak SNR with a SNR ˃ 1000:1, ˃500:1, ˃ and The CT-cHSQC experiment (46) was taken with 50 and 17408 points, a spectral window of 1680.3 Hz (2.4 ppm) and 8802.8 Hz (50 ppm), and a carrier frequency of 4.2 ppm and 83 ppm in ^1^H and ^13^C, respectively. Data were zero-filled to 2X the total points collected in ^1^H and ^13^C and a cosine squared function was applied in both dimensions. The CT-cHSQC experiment was processed with SMILE-based linear prediction (35).

### Transglycosidic Torsion Calculations

Transglycosidic ^3^*J*_CH_ values obtained from the PIP-HSQMBC experiment were substituted into Zhang et al. (39) and Reeves and Wang (40) parameterized Karplus relations in order to calculate φ and ψ torsions. These were:

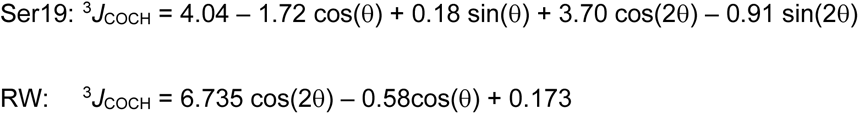

To account for the uncertainty introduced by using these equations, the error was determined by taking the square root of experimental error squared plus the Karplus-like RMSE squared.

The above Karplus-like equations were used to obtain torsion values from experimental ^3^*J*_CH_ values. The converse was also utilized, namely, we predicted ^3^*J*_CH_ values from the MD trajectories. While Ser19 can be directly used for this goal, Reeves and Wang proposed a related equation in the same work to use with MD data which is the one used in the present work, that considered in addition to the torsion angle the central C-O distance (*r*_CO_) from the MD simulation, expressed by the following equation:

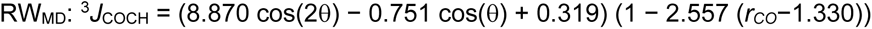

### Molecular dynamics simulations

MD simulations were performed using the AMBER 20 software package (47) on a workstation equipped with four GeForce GTX 1080Ti Graphic Processing Units (GPU) for oligomers up to 10 residues. For the larger oligomers (up to 12 GXM repeating units) simulations were run at the FDA core facility on NVIDIA A100 GPU. All structure and parameter files for pentamannose, GXM10 w/wo Ac_3_ and GXM-12RU were created using the TLEaP module included in AMBER with the force field Glycam-06j-1 (48). Following analysis of initial MD runs it became apparent that the xylose rings needed reinforcement to prevent intermittent puckering exchange from the normal ^4^C_1_ chair to the ^1^C_4_ form. This was achieved by using parabolic restraints on two torsion angles in xylose defined to atoms [O5-C1-C2-C3] and [C3-C4-C5-O5]. Analysis was performed on these torsions to verify the effect of restraints was limited to prevent puckering changes.

All initial conformers were solvated with TIP3FB explicit water models (49) defining cubic cells for periodic conditions with a minimum distance from the oligosaccharide to the border of 8 Å, except for GXM-12RU for which an octahedral cell was employed to reduce the number of water molecules needed. Na^+^ counterions were used for neutralization of charges. For GXM-12RU the cell linear dimensions were of ~170 Å containing in addition to the PS 12 sodium ions and over 79,000 waters. Non-bonded van der Waals and electrostatic scaling factors for 1-4 interactions were set to unity (SCEE = SCNB = 1) as required by Glycam. Long range electrostatic interactions were computed with particle-mesh Ewald summation (50), with a non-bonded cutoff distance of 8 Å. Starting oligomer conformations with default torsion values were appropriate to proceed with energy minimization in all cases. Then, a 1 ns preparation MD run was used to equilibrate the system to the target temperature of 300K for production, as follows: 0K to 310K in 0.1 ns, 310 K from 0.1 to 0.6 ns, 310K to 300K from 0.6 to 0.8 ns, 300K from 0.8 to 1.0 ns. MD target durations for analysis were of 2 μs at 300K, with NPT conditions and hydrogen mass repartition (HMR) (51) to allow an integration time for the equations of motion of 4 fs, with hydrogen-containing covalent bonds constrained to their equilibrium lengths using the SHAKE algorithm (52). As mentioned in the text, for GXM10-Ac_3_ the first 100 ns were regarded as equilibration based on 2D RMSD analysis, Figure S6.

Processing of trajectories was performed using *cpptraj*, included in AMBER, and python in-house scripts. General MD statistical analysis on GXM10-Ac_3_ was performed over 2 μs utilizing 50,000 frames (1 every 40 ps). For distance analysis and NOE predictions we utilized the ‘type noe’ option in *cpptraj* that reports the average distance <*r*^−6^>^−1/6^. Free energy torsion maps were produced from bin populations using *G_i_ = −k_B_T* ln(*N_i_/N_Max_*), where *k_B_* is the Boltzmann’s constant, *T* is the MD temperature, *N_i_* is the population of bin *i* and *N_Max_* is the population of the most populated bin. ^3^*J*_CH_-couplings were calculated using the equations described in the previous section from instantaneous torsion values, then standard statistics definitions were used to obtain the average and standard deviation values for each case. Figures were created with OriginPro 2019 (www.OriginLab.com; Northampton, MA), and PyMol (www.schrodinger.com/pymol). Movies were generated with PyMol.

## Acknowledgments

C.J.C was funded by an Irish Research Councill Postgraduate award (GOIPG/2016/998). S.O by SFI 13/IA/1959 and 20/FFP-P/884. A.C. is supported in part by National Institutes of Health grants AI052733, AI152078, and HL059842. M.P.W. is supported by NIH grants AI162381 and AI152078. H.F.A. thanks Xiaocong Wang and Rob Woods for helpful advice to restraint xylose rings. A.A.H. thanks Jeahoo Kwon for help setting up initial NMR experiments and writing a program to aid in torsion calculations from NMR data.

## Notes

### Competing Interest Statement

The authors have declared no competing interest.

