## Supplemental Information for "The structure of a *C. neoformans* polysaccharide motif recognized by protective antibodies: A combined NMR and MD study"

**This PDF file includes:**
Figures S1 to S8
Tables S1 to S4
Supplemental movie legend SM1 to SM3

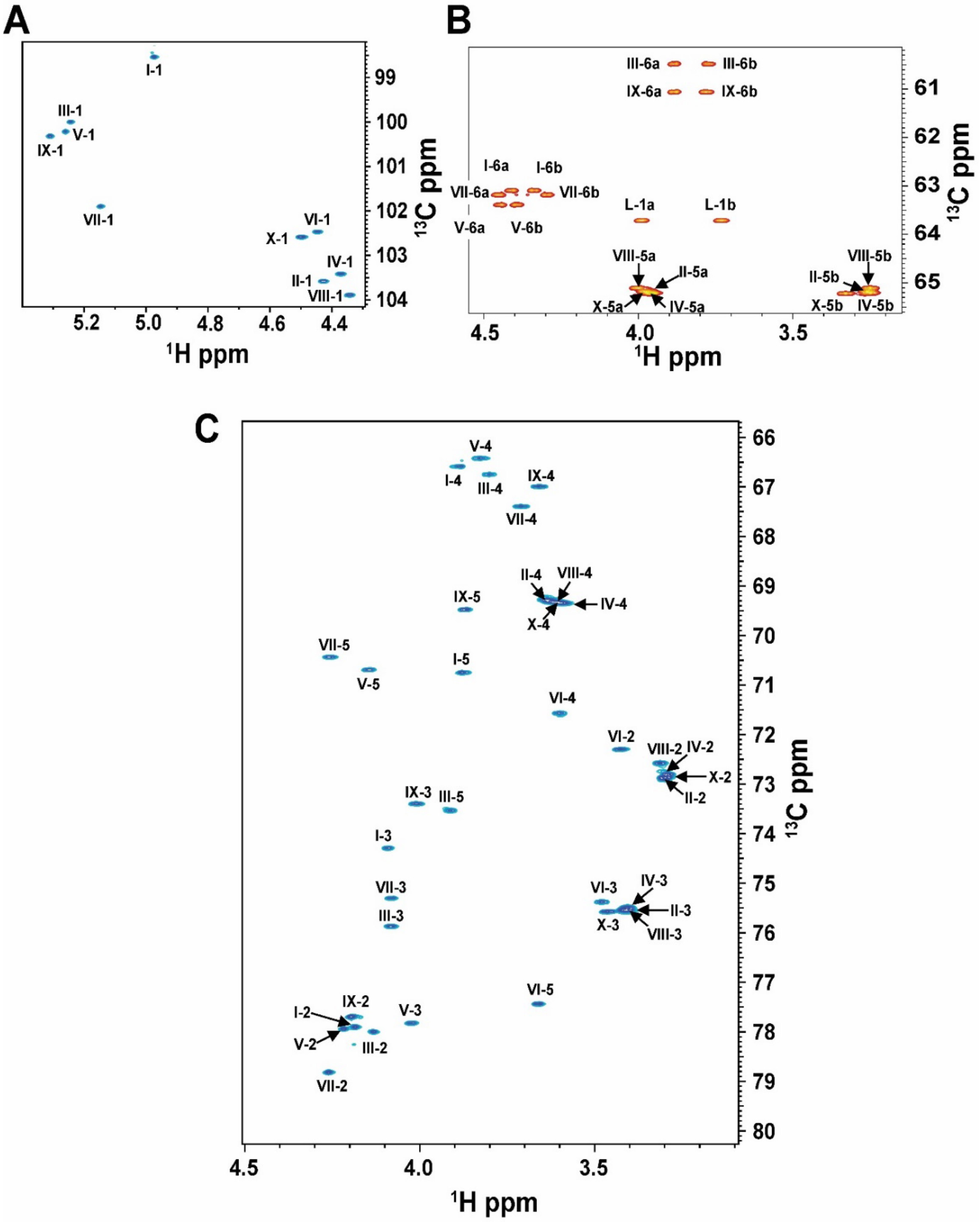


**Figure S1: GXM GXM10-Ac$_3$ $^{1}$H-$^{13}$C eHSQC. (A)** GXM10-Ac$_3$ Anomeric ($^{1}$H, $^{13}$C) region have a unique chemical shift ($^{1}$H: 4.3 – 5.4 ppm; $^{13}$C: 98-104 ppm) dispersed from other ring $^{1}$Hs and $^{13}$Cs. **(B)** In the eHSQC experiment, CH$_2$ resonances are negative, and *O*-acetylated Man C6s (Man[I,V,VII]) are de-shielded from the Man C6s without *O*-acetylation (Man[II, IX]) **(C)** NMR signals for Xyl rings are heavily overlapped, particularly for H2C2, H3C3, H4C4, and H5C5.

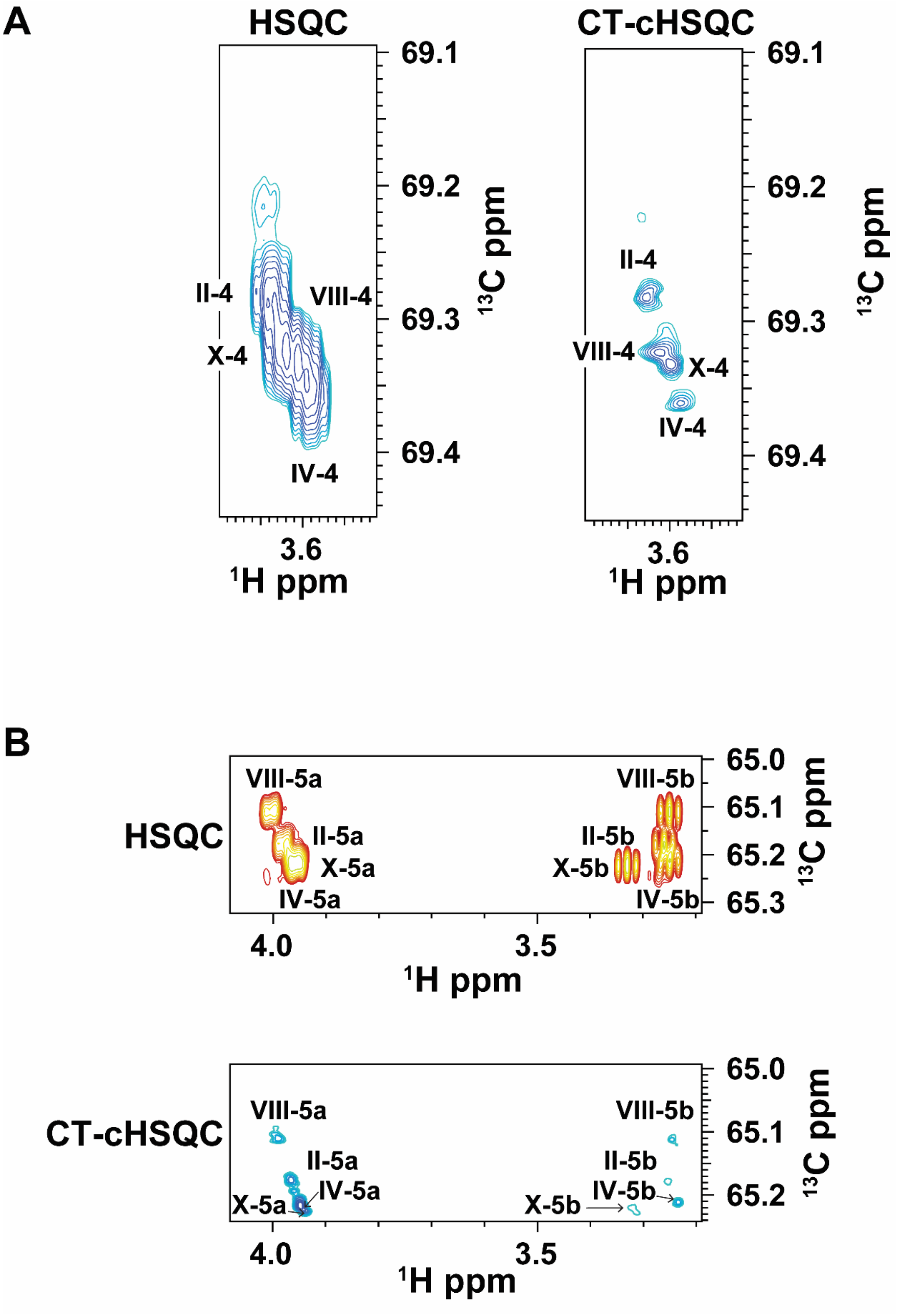


**Figure S2: Overlapping H4C4 (A) and H5C5 (B) Xylose resonances in the $^{1}$H-$^{13}$C HSQC experiment are resolved in a $^{1}$H-$^{13}$C CT-cHSQC experiment.**

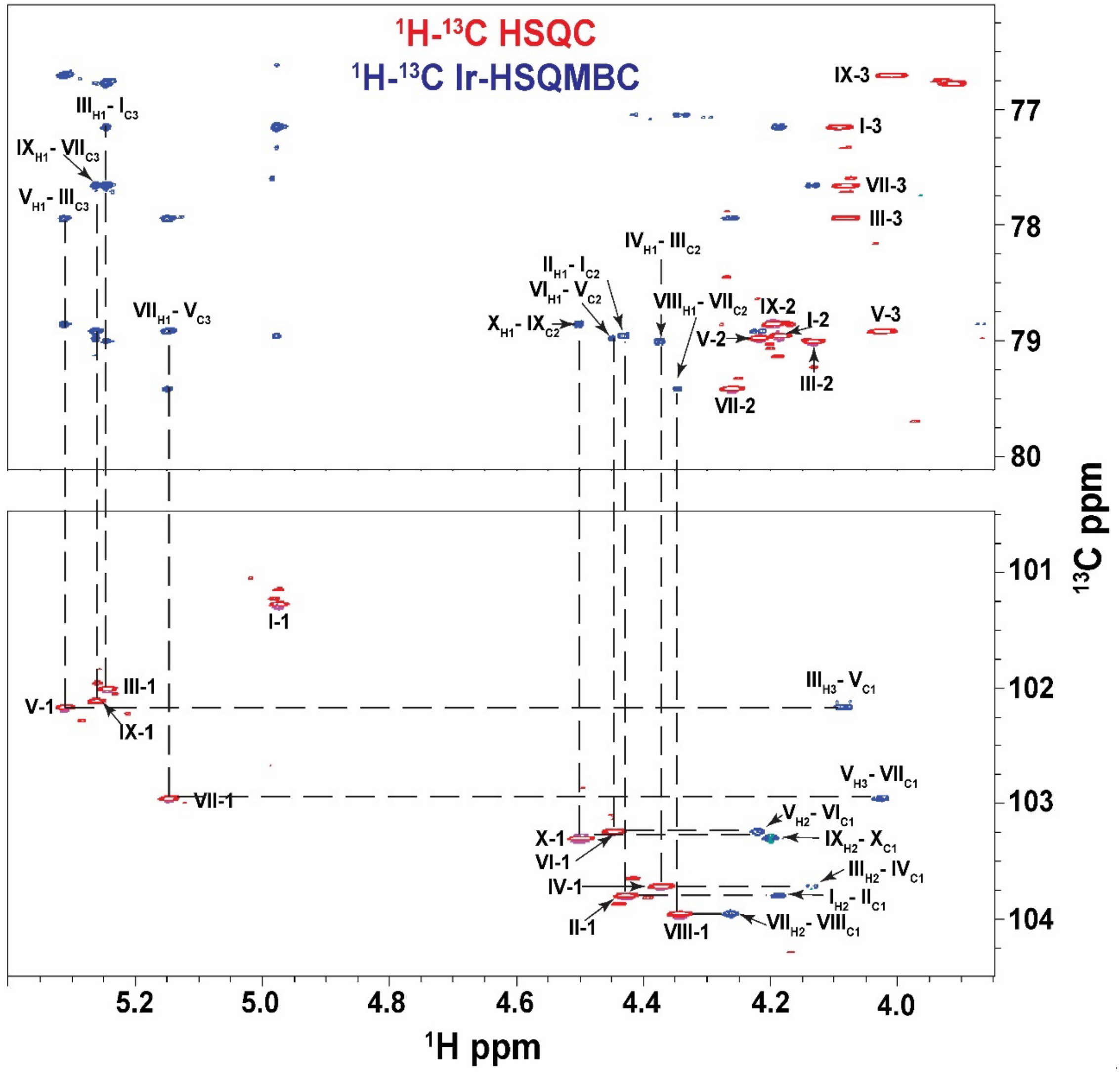


**Figure S3: GXM10-Ac$_3$ inter-residue linkages were assigned from inter-glycan crosspeaks obtained in a $^1$H-$^{13}$C LR-HSQMBC experiment (blue). $^1$H-$^{13}$C HSQC spectrum is overlayed in red.**

**Table S1: GXM Synthetic Decasaccharide $^{1}H$-$^{13}C$ Chemical Shift Assignments**

| | | I | II | III | IV | V | VI | VII | VIII | IX | X |
|---|---|---|---|---|---|---|---|---|---|---|---|
| $^{1}H$ | **H-1** | 4.97 | 4.43 | 5.24 | 4.37 | 5.26 | 4.45 | 5.15 | 4.34 | 5.31 | 4.50 |
| | **H-2** | 4.19 | 3.31 | 4.13 | 3.29 | 4.22 | 3.43 | 4.26 | 3.32 | 4.20 | 3.30 |
| | **H-3** | 4.09 | 3.42 | 4.08 | 3.41 | 4.03 | 3.48 | 4.08 | 3.41 | 4.01 | 3.47 |
| | **H-4** | 3.89 | 3.69 | 3.80 | 3.60 | 3.83 | 3.60 | 3.71 | 3.64 | 3.66 | 3.63 |
| | **H-5ax** | 3.88 | 3.27 | 3.91 | 3.26 | 4.15 | 3.66 | 4.26 | 3.25 | 3.87 | 3.33 |
| | **H-5eq** | - | 3.99 | - | 3.97 | - | - | - | 4.00 | - | 3.96 |
| | **H-6a** | 4.41 | - | 3.89 | - | 4.45 | - | 4.45 | - | 3.89 | - |
| | **H-6b** | 4.34 | - | 3.77 | - | 4.39 | - | 4.30 | - | 3.78 | - |
| | **H-$CH_3$** | 2.14 | - | - | - | 2.20 | - | 2.17 | - | - | - |
| $^{13}C$ | **C-1** | 98.54 | 103.58 | 100.00 | 103.58 | 100.22 | 102.48 | 101.90 | 103.89 | 100.32 | 102.59 |
| | **C-2** | 77.91 | 72.89 | 78.00 | 72.89 | 77.94 | 72.30 | 78.82 | 72.58 | 77.70 | 72.87 |
| | **C-3** | 74.29 | 75.53 | 75.31 | 75.53 | 77.83 | 75.38 | 75.87 | 75.55 | 73.40 | 75.57 |
| | **C-4** | 66.59 | 69.36 | 66.75 | 69.36 | 66.42 | 71.57 | 67.40 | 69.28 | 67.00 | 69.32 |
| | **C-5** | 70.75 | 65.19 | 73.53 | 65.19 | 70.69 | 77.44 | 70.44 | 65.11 | 69.48 | 65.22 |
| | **C-6** | 63.10 | - | 60.49 | - | 65.39 | | 63.19 | - | 61.06 | - |
| | **C-CO** | 174.08 | - | - | - | 174.16 | - | 174.12 | - | - | - |
| | **C-$CH_3$** | 20.24 | - | - | - | 20.58 | - | 20.57 | - | - | - |

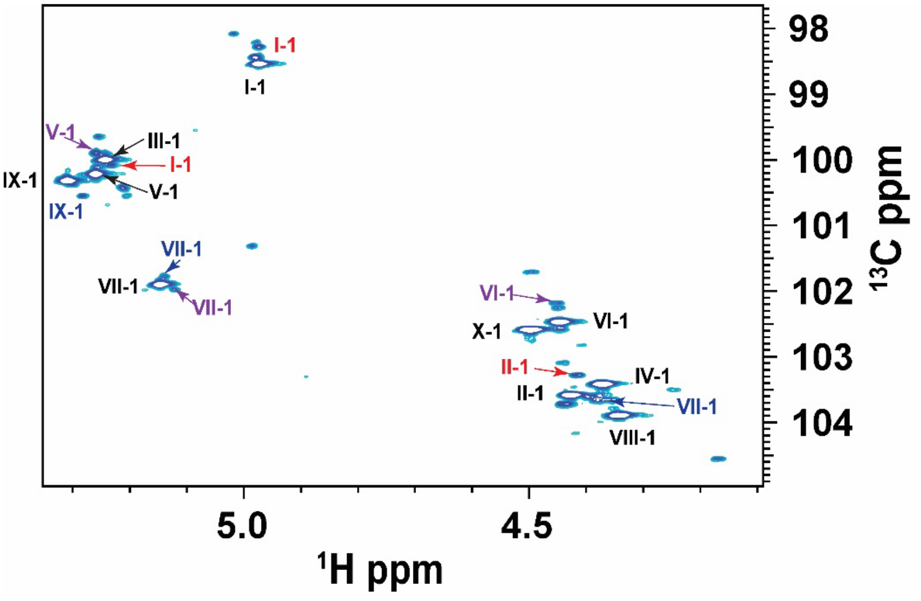


**Figure S4: $^{1}$H-$^{13}$C HSQC Anomeric Region showing de-*O*-Acetylated Decasaccharide Resonances.** The black labels designate GXM10-Ac3 anomeric signals. The loss of *O*-acetylation at I-Man-C6, V-Man-C6, and VII-Man-C6 are labeled in red, purple, and blue, respectively.

**Table S2 $^{1}H$-$^{13}C$ Chemical Shift Assignments for de-O-Acetylated GXM Decasaccaharide**

| | | I | II | III | IV | V | VI | VII | VIII | IX | X |
|---|---|---|---|---|---|---|---|---|---|---|---|
| $^{1}H$ | H-1 | *4.97* | *4.42* | *5.23* | 4.37 | 5.26* | 4.45* | 5.12*(5.14) | 4.34 | (5.28) | 4.50 |
| | H-2 | *4.17* | 3.31 | 4.13 | 3.29 | 4.22 | 3.43 | 4.26 | 3.32 | 4.20 | 3.30 |
| | H-3 | *4.08* | 3.42 | *4.08* | 3.41 | 4.03 | 3.48 | 4.08 | 3.41 | 4.01 | 3.47 |
| | H-4 | *3.85* | 3.69 | 3.80 | 3.60 | 3.83 | 3.60 | 3.71 | 3.64 | 3.66 | 3.63 |
| | H-5ax | 3.88 | 3.27 | *3.91* | 3.26 | 4.15 | 3.66 | 4.26 | 3.25 | 3.87 | 3.33 |
| | H-5eq | - | 3.99 | - | 3.97 | - | - | - | 4.00 | - | 3.96 |
| | H-6a | 4.41 | - | 3.89 | - | 4.45 | - | 4.45 | - | 3.89 | - |
| | H-6b | 4.34 | - | 3.77 | - | 4.39 | - | 4.30 | - | 3.78 | - |
| | H-$CH_3$ | 2.14 | - | - | - | 2.20 | - | 2.17 | - | - | - |
| $^{13}C$ | C-1 | *98.28* | *103.28* | *100.08* | 103.58 | 99.65* | 102.18* | 101.98*(101.78) | 103.89 | (100.55) | 102.59 |
| | C-2 | *77.72* | 72.89 | 78.00 | 72.89 | 77.94 | 72.30 | 78.82 | 72.58 | 77.70 | 72.87 |
| | C-3 | *74.65* | 75.53 | *75.41* | 75.53 | 77.83 | 75.38 | 75.87 | 75.55 | 73.40 | 75.57 |
| | C-4 | 66.59 | 69.36 | 66.75 | 69.36 | 66.42 | 71.57 | 67.40 | 69.28 | 67.00 | 69.32 |
| | C-5 | *73.23* | 65.19 | *73.50* | 65.19 | 70.69 | 77.44 | 70.44 | 65.11 | 69.48 | 65.22 |
| | C-6 | 63.10 | - | 60.49 | - | 65.39 | | 63.19 | - | 61.06 | - |
| | C-CO | 174.08 | - | - | - | 174.16 | - | 174.12 | - | - | - |
| | C-$CH_3$ | - | - | - | - | - | - | - | - | - | - |

Italic denotes chemical shift from the loss of I-Man-C6 O-Acetylation, * denotes the chemcial shift from the loss of V-Man-C6 O-Acetylation, and () denotes loss of VII-Man-C6 O-Acetylation.

**Table S3. Measured transglycosidic $^3J_{CH}$ values** from $^1$H-$^{13}$C PIP-HSQMBC NMR experiment. All $^3J_{CH}$ have an error of 0.5 Hz. $\phi$ and $\psi$ torsions were calculated using two different parameterized Karplus relations (see methods). Gold color denotes torsions similar to the average torsions obtained from the MD trajectory.

| Tglyc. Bond | | $^3J_{CH}$ Exp (Hz) | Calc. Torsion from $J_{meas.}$ | | | | | |
|---|---|---|---|---|---|---|---|---|
| | | | Tor-S19 | | | | Tor-RW | |
| Man[I]-Man[III] | $\phi$ | 2.5 | -123 (+/- 7) | -55 (+/- 8) | 40 (+/- 8) | 110 (+/- 7) | ± 51 (+/- 12) | ± 124 (+/- 11) |
| | $\psi$ | -- | -- | -- | -- | -- | -- | -- |
| Man[III]-Man[V] | $\phi$ | 3.0 | -126 (+/- 7) | -51 (+/- 8) | 37 (+/- 8) | 113 (+/- 6) | ± 47 (+/- 11) | ± 127 (+/- 10) |
| | $\psi$ | -- | -- | -- | -- | -- | -- | -- |
| Man[V]-Man[VII] | $\phi$ | 3.5 | -130 (+/- 6) | -46 (+/- 8) | 32 (+/- 6) | 116 (+/- 6) | ± 42 (+/- 11) | ± 131 (+/- 10) |
| | $\psi$ | 3.4 | -129 (+/- 6) | -47 (+/- 8) | 33 (+/- 8) | 116 (+/- 6) | ± 43 (+/- 11) | ± 131 (+/- 10) |
| Man[VII]-Man[IX] | $\phi$ | 2.9 | -126 (+/- 7) | -52 (+/- 8) | 38 (+/- 8) | 112 (+/- 7) | ± 48 (+/- 11) | ± 126 (+/- 10) |
| | $\psi$ | 4.1 | -134 (+/- 6) | -40 (+/- 9) | 26 (+/- 9) | 121 (+/- 6) | ± 36 (+/- 12) | ± 136 (+/- 10) |
| Man[I]-Xyl[II] | $\phi$ | 3.8 | -132 (+/- 6) | -44 (+/- 8) | 29 (+/- 8) | 118 (+/- 6) | ± 40 (+/- 11) | ± 134 (+/- 10) |
| | $\psi$ | 5.5 | -143 (+/- 6) | -25 (+/- 14) | 11 (+/- 14) | 130 (+/- 6) | ± 22 (+/- 17) | ± 148 (+/- 11) |
| Man[III]-Xyl[IV] | $\phi$ | 4.0 | -134 (+/- 6) | -41 (+/- 9) | 27 (+/- 9) | 121 (+/- 6) | ± 37 (+/- 12) | ± 136 (+/- 10) |
| | $\psi$ | 5.2 | -142 (+/- 6) | -28 (+/- 16) | 14 (+/- 16) | 128 (+/- 6) | ± 25 (+/- 19) | ± 146 (+/- 11) |
| Man[V]-GlcA[VI] | $\phi$ | 4.1 | -134 (+/- 6) | -41 (+/- 9) | 27 (+/- 9) | 121 (+/- 6) | ± 37 (+/- 12) | ± 136 (+/- 10) |
| | $\psi$ | 4.9 | -139 (+/- 6) | -33 (+/- 11) | 19 (+/- 11) | 126 (+/- 6) | ± 29 (+/- 14) | ± 143 (+/- 5) |
| Man[VII]-Xyl[VIII] | $\phi$ | 4.2 | -135 (+/- 6) | -40 (+/- 9) | 25 (+/- 9) | 121 (+/- 6) | ± 36 (+/- 12) | ± 137 (+/- 10) |
| | $\psi$ | 4.8 | -139 (+/- 6) | -34 (+/- 10) | 20 (+/- 10) | 125 (+/- 6) | ± 30 (+/- 13) | ± 142 (+/- 10) |
| Man[IX]-Xyl[X] | $\phi$ | 3.5 | -130 (+/- 6) | -46 (+/- 8) | 32 (+/- 8) | 117 (+/- 6) | ± 42 (+/- 11) | ± 131 (+/- 10) |
| | $\psi$ | 4.8 | -140 (+/- 6) | -33 (+/- 10) | 19 (+/- 10) | 125 (+/- 6) | ± 29 (+/- 14) | ± 142 (+/- 10) |

**Table S4. Average ϕ and ψ torsions from MD trajectory and predicted $^3J_{CH}$ values.** Gray denotes calculated $^3J_{CH}$ values that do not agree well with experimental torsions.

| Tglyc. Bond | | MD Tor. | $^3J_{CH}$ Calc. from MD | |
|---|---|---|---|---|
| | | | *J*-S19 | *J*-RW |
| Man[I]-Man[III] | ϕ | -53 (+/- 12) | 2.7 (+/-1.1) | 2.0 (+/-1.0) |
| | ψ | -8 (+/- 21) | *5.3* (+/-0.9) | *4.3* (+/-1.0) |
| Man[III]-Man[V] | ϕ | -51 (+/- 14) | 2.9 (+/-1.1) | 2.2 (+/-1.0) |
| | ψ | -5 (+/- 22) | *5.3* (+/-1.0) | *4.3* (+/-1.0) |
| Man[V]-Man[VII] | ϕ | -45 (+/- 15) | 3.4 (+/-1.1) | 2.6 (+/-1.0) |
| | ψ | 5 (+/- 24) | *4.9* (+/-1.3) | *4.2* (+/-1.1) |
| Man[VII]-Man[IX] | ϕ | -47 (+/- 14) | 3.3 (+/-1.1) | 2.5 (+/-1.0) |
| | ψ | 6 (+/- 22) | *5.0* (+/-1.2) | *4.3* (+/-1.0) |
| Man[I]-Xyl[II] | ϕ | 41 (+/- 20) | 2.4 (+/-1.5) | 2.8 (+/-1.3) |
| | ψ | 14 (+/- 25) | 4.7 (+/-1.7) | 4.0 (+/-1.3) |
| Man[III]-Xyl[IV] | ϕ | 41 (+/- 31) | 2.6 (+/-1.8) | 2.9 (+/-1.3) |
| | ψ | 17 (+/- 26) | 4.3 (+/-1.8) | 3.8 (+/-1.4) |
| Man[V]-GlcA[VI] | ϕ | 39 (+/- 30) | 2.7 (+/-1.9) | 3.0 (+/-1.4) |
| | ψ | 32 (+/- 33) | 2.9 (+/-1.9) | 2.8 (+/-1.5) |
| Man[VII]-Xyl[VIII] | ϕ | 40 (+/- 23) | 2.6 (+/-1.6) | 3.0 (+/-1.3) |
| | ψ | 30 (+/- 27) | 3.4 (+/-2.0) | 3.2 (+/-1.5) |
| Man[IX]-Xyl[X] | ϕ | 40 (+/- 22) | 2.4 (+/-1.4) | 2.8 (+/-1.2) |
| | ψ | 20 (+/- 30) | 4.0 (+/-1.8) | 3.5 (+/-1.3) |

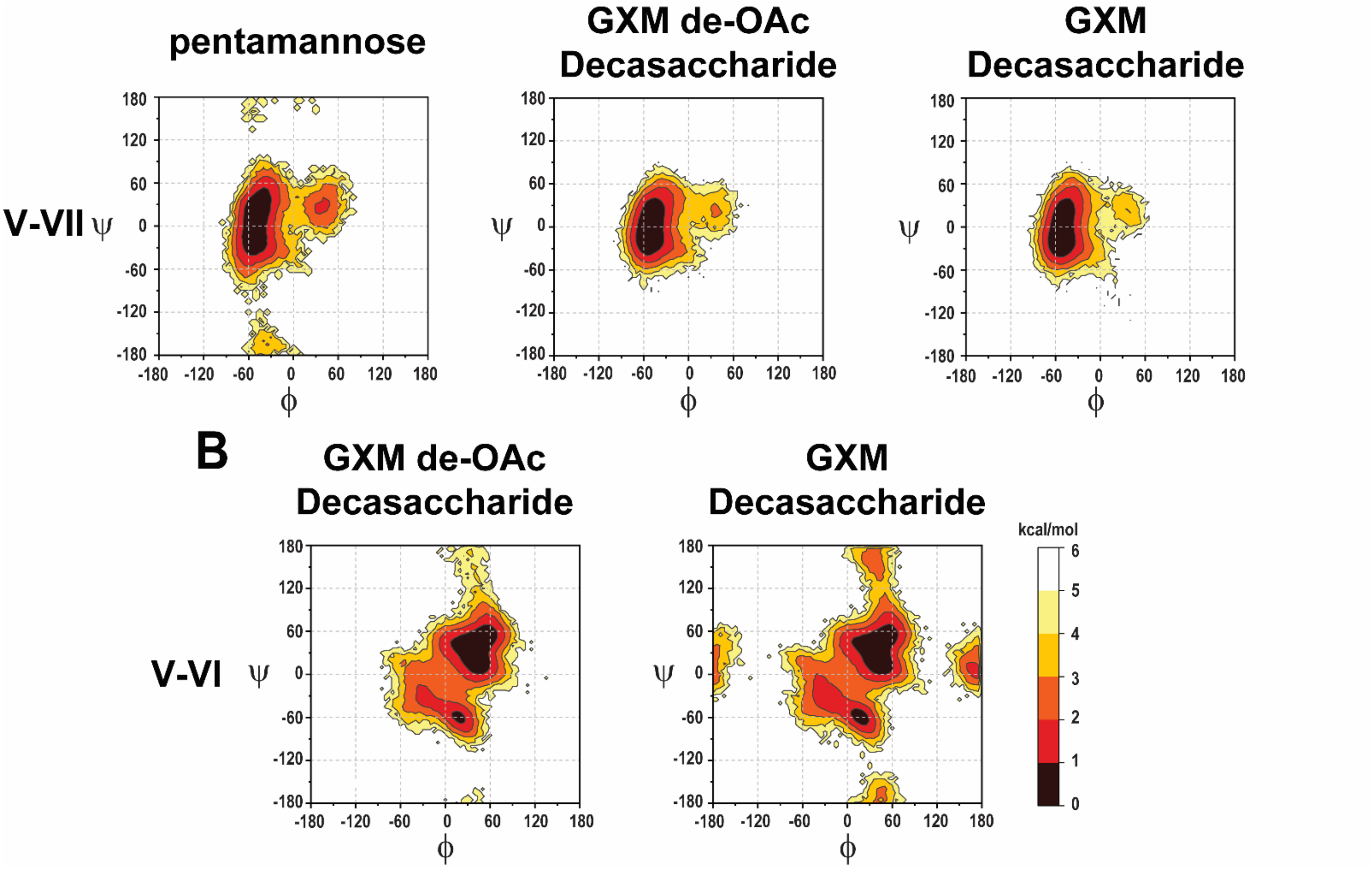


**Figure S5: Effect of Substitution on GXM10-Ac$_3$ Structure.** A ($\phi$, $\psi$) transglycosidic torsion angle population analysis of a pentamannose, GXM10, and GXM10-Ac$_3$ were compared. **(A)** The Man[V]-Man[VII] transglycosidic torsion heatmap suggests a decrease in flexibility upon substitution of Xyl and *O*-acetylation. **(B)** *O*-acetylaton does not seem to have an effect on the torsional landscape of the Man-Xyl/GlcA linkages.

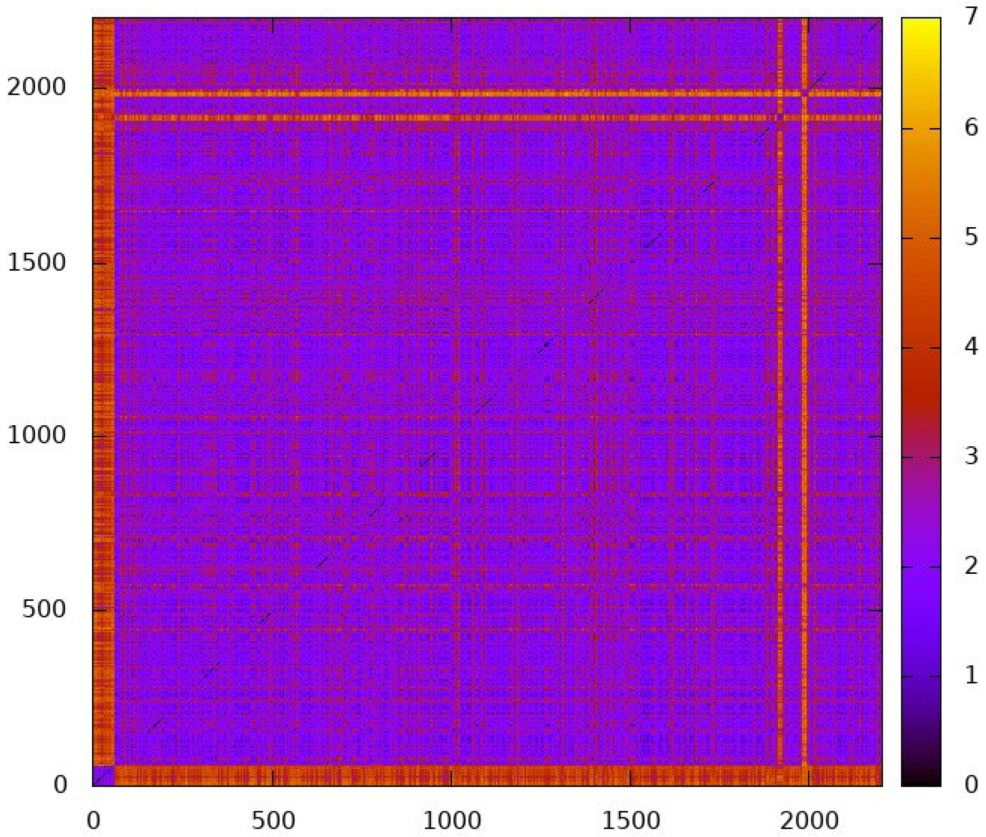


.

**Figure S6.** 2D-RMSD map of GXM10-$Ac_3$, analyzing 2200 frames over 2.2 μs MD. RMSD calculated using the 6-ring atoms for all 10 glycan residues in GXM10-$Ac_3$ (total of 60 atoms). The first 75 ns correspond to a high energy conformation that is not revisited once a steady state is reached. Based on this, analysis of the GXM10-$Ac_3$ MD was limited the resulting trajectory starting at 100 ns up to 2100 ns (2 μs in total)

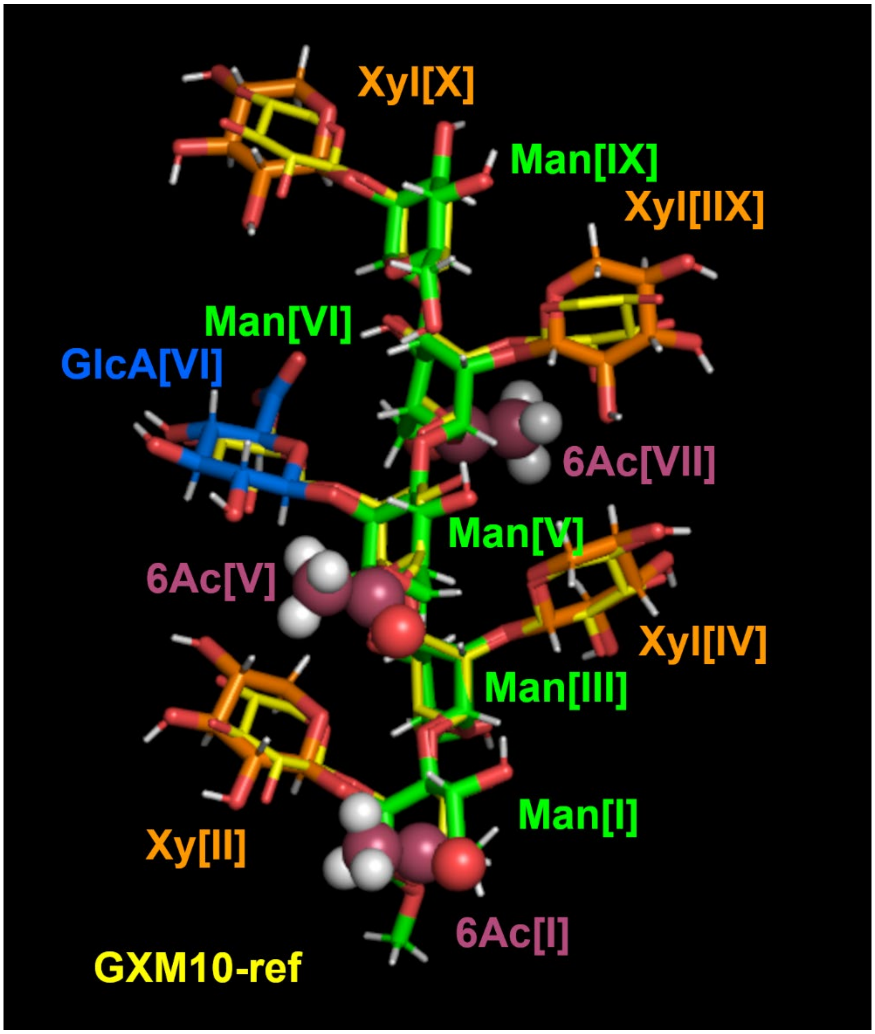


**Figure S7.** Overlay of GXM10ref (yellow) and the GXM10-$Ac_3$ model with lowest RMSD to GXM10ref (RMSD = 0.56 Å, 60 ring atoms) from the 2 μs MD trajectory.

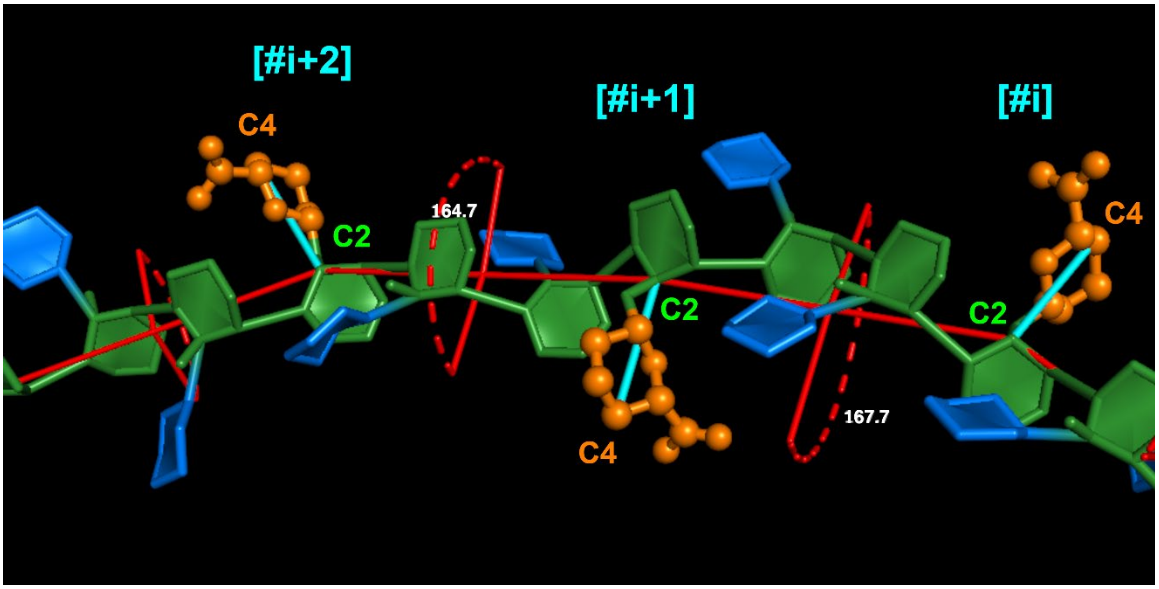


**Figure S8. Vector definitions to calculate the twist pitch of the GXM PS.** Vectors between atoms Man{GlcA}-C2 and GlcA-C4 were defined for each repeating unit (RU) [cyan lines], then dihedral angles between consecutive vectors were calculated throughout the trajectory. The figure shows a portion of the final frame of GXM12RU to illustrate the general procedure employed to determine inter-RU's dihedrals to generate Figure 4C.

**Movie S1.** GXM10-ref shown in Figure 3A rolled around the *y*-axis to better appreciated its 3D structure.

**Movie S2.** The trajectory of the linked Man[V]6Ac-GlcA shown in Figure 3B, to better illustrate the branch dynamics. The trajectory was ‘smoothed’ to reduce high frequency movements.

**Movie S3.** Last 500 ns of the MD trajectory of GXM12RU. Frames were RMSD minimized to the ring atoms of the colored residues. The trajectory was ‘smoothed’ to reduce high frequency movements.
